# Genome-wide mapping of histone modifications in two species of *Leptosphaeria maculans* showing contrasting genomic organization and host specialization

**DOI:** 10.1101/2020.05.08.084566

**Authors:** J.L. Soyer, C. Clairet, E.J. Gay, N. Lapalu, T. Rouxel, E.H. Stukenbrock, I. Fudal

## Abstract

In plant-associated fungi, the role of the epigenome is increasingly recognized as an important regulator of genome structure and of the expression of genes involved in interaction(s) with the host plant. Two closely-related phytopathogenic species, *Leptosphaeria maculans* ‘brassicae’ (Lmb) and *L. maculans* ‘lepidii’ (Lml) exhibit a large conservation of genome synteny but contrasting genome structure. Lmb has undergone massive invasion of its genome by transposable elements amounting to one third of its genome and clustered in large TE-rich regions on chromosomal arms, while Lml genome has only a small amount of repeats (3% of the genome). Previous studies showed that the TE-rich regions of Lmb harbour a few species-specific effector genes, expressed during plant infection. The distinct genome structures shown by Lmb and Lml thus provides an excellent model for comparing the organization of pathogenicity/effector genes in relation to the chromatin landscape in two closely related phytopathogenic fungi. Here, we performed chromatin immunoprecipitation during axenic culture, targeting either histone modifications typical for heterochromatin or euchromatin, combined with transcriptomic analysis to analyse the influence of chromatin organisation on gene expression. In both species, we found that facultative heterochromatin landscapes associated with H3K27me3-domains are enriched with genes lacking functional annotation, including numerous candidate effector and species-specific genes. Notably, orthologous genes located in H3K27me3-domains in both species are enriched with genes encoding putative proteinaceous and metabolic effectors. These genes are mostly silenced in axenic growth conditions and are likely to be involved in interaction with the host. Compared to other fungal species, including Lml, Lmb is distinct in having large H3K9me3-domains associated with TE-rich regions that contain numerous species-specific effector-encoding genes. Discovery of these two distinctive heterochromatin landscapes now raises questions about their involvement in the regulation of pathogenicity, the dynamics of these domains during plant infection, and the selective advantage to the fungus to host effector genes in H3K9me3- or H3K27me3-domains.

## Introduction

Each year, hundreds of millions of tons of agricultural crops are devastated by plant pathogenic fungi or made unfit for consumption due to contamination by mycotoxins (Fisher *et al*., 2018). Management strategies to control fungal infection mainly involve chemical control or breeding for naturally resistant crop cultivars. However, fungal plant pathogens have proven capable of rapidly evolving resistance against implemented fungicides (Fisher *et al*., 2018) and to overcome specific plant resistance genes within a few years (for instance in *Leptosphaeria maculans*; Rouxel and Balesdent, 2017), emphasising the need for improved control methods.

Understanding the determinants of the extreme adaptive abilities of fungal plant pathogens is a critical issue for the development of effective and sustainable control methods. In that respect, comparative and population genomic analyses have provided new insights into the evolutionary dynamics of fungal plant pathogens (e.g. Stukenbrock *et al.*, 2011; Grandaubert *et al.*, 2014; Sánchez-Vallet *et al.*, 2018). Notably, transposable elements (TE) have been shown to play a crucial role in shaping the genome structure of plant-pathogenic fungi. TEs are often organized in clusters, compartmentalizing the genome into gene-rich regions and TE-rich regions (e.g. in *L. maculans* and in *Mycosphaerella fijiensis*; Rouxel *et al.*, 2011; Ohm *et al.*, 2012; Grandaubert *et al*., 2014). While the gene density in TE-rich regions is low, genes located in these regions have been shown to evolve faster than genes located in TE-poor regions (Croll & McDonald, 2012; de Jonge *et al.*, 2013). Interestingly, rapidly evolving genes in TE-rich regions have, in many cases, been identified as genes involved in niche adaptation and notably effector genes. Effectors are considered as key elements of pathogenesis, allowing pathogens to circumvent host recognition, impede defence reactions and facilitate host invasion. Effectors are mainly secreted proteins, but can also correspond to secondary metabolites and small RNAs (Lo Presti *et al*., 2015; Collemare *et al*., 2019; Rocafort *et al*., 2020). Plants have evolved strategies to recognize and counteract effectors, exposing them to a strong selection pressure by the host immune system (Stergiopoulos *et al*., 2007; Chuma *et al*., 2011; Rouxel and Balesdent, 2017). Indeed, in the course of the co-evolution between a pathogen and its host, the host has developed an active immune system allowing the direct or indirect recognition of some effector molecules, to activate defence responses, often involving a local cell death, called the hypersensitive response. Effectors that can be recognized by the host are called avirulence proteins (Jones and Dangl, 2006; Lo Presti *et al*., 2015).

*Leptosphaeria maculans* ‘brassicae’ (hereinafter referred to as Lmb) belongs to the Dothideomycete class of Ascomycete fungi and is responsible for causing stem canker of oilseed rape (*Brassica napus*). Lmb displays a complex, hemibiotrophic life cycle, during which it alternates between different nutritional modes on its host plant. It causes necrosis on different plant organs: leaves, more rarely seedpods, and the base of the stem, causing lodging of the plant and yield losses (Rouxel and Balesdent, 2005). The most efficient method of disease control relies on the use of major resistance genes present in oilseed rape and other Brassica species. Although efficient, this control method is not sustainable, as Lmb is able to “break down” novel sources of genetic resistance rapidly (Rouxel and Balesdent, 2017). One third of the Lmb genome is made of TE-rich regions. These regions are enriched with putative effector genes and include all currently known avirulence genes. These avirulence genes are highly expressed in the first seven days of leaf and cotyledon infection (Gout *et al.*, 2006; Fudal *et al.*, 2007; Parlange *et al.*, 2009; Balesdent *et al.*, 2013; van de Wouw *et al*., 2014; Ghanbarnia *et al*., 2015; 2018; Plissonneau *et al.*, 2016; 2018; Petit-Houdenot *et al.*, 2019). Another set of putative proteinaceous effector genes are located in gene-rich regions of the genome and are specifically expressed during stem infection (Gervais *et al*., 2017).

In eukaryotic cells, chromatin can adopt different conformational states directly influencing gene expression: gene-rich euchromatin, sheltering constitutively expressed genes, and gene-poor heterochromatin, in which genes are silent. The different chromatin states are characterized by different post-translational modifications of histones around which DNA is wrapped. Typically, heterochromatin is enriched in the trimethylation of the lysine 9 of histone H3 (H3K9me3) and lysine 27 (H3K27me3) while euchromatin is enriched in the di- (or tri-) methylation of the lysine 4 of histone H3 (H3K4me2) (Zhao and Garcia, 2015). Recent research has focused on dynamic changes in DNA accessibility and how these may play fundamental roles in the adaptation of different individuals to abiotic and biotic stresses (Gomes and Pelosi, 2013; Zogli and Libault, 2017). In fungal plant pathogens or endophytes, evidence is accumulating that transcriptional reprogramming of effector genes (either proteinaceous or metabolic) is tightly controlled by chromatin-based regulatory mechanisms (for example in *Fusarium graminearum, Epichloe festucae*, Lmb and *Zymoseptoria tritici*; Connolly *et al*., 2013; Chujo and Scott, 2014; Soyer *et al*., 2014; Soyer *et al*., 2015a; Soyer *et al*., 2019). In Lmb, the location of avirulence genes in TE-rich regions has an influence on their evolution under selection pressure (Rouxel *et al*., 2011; Daverdin *et al*., 2012) and plays a role in regulating their expression during axenic culture via deposition of H3K9me3 (Soyer *et al*., 2014). Lmb belongs to the *L. maculans* / *Leptosphaeria biglobosa* species complex, comprising species with different host specialization and genome organization (Grandaubert *et al.*, 2014). Within the species complex, the *L. maculans* species infecting oilseed rape, Lmb, is the only one having large regions of its genome enriched with TEs while other genomes have a low TE-content (∼3-15% compared to more than 30% TEs in Lmb; Rouxel *et al*., 2011; Grandaubert *et al*., 2014; Dutreux *et al*., 2018). For example, the genome of the species that is most closely related to Lmb, *Leptosphaeria maculans* ‘lepidii’ (hereinafter referred to as Lml), which infects crucifers such as *Lepidium sativum*, and to a lesser extent *Camelina sativa* and *Brassica rapa* (Petrie, 1969), has a low TE content (3% of TEs; Grandaubert *et al*., 2014). Interestingly, the genomes of Lmb and Lml show a high level of macrosynteny, with only a few intra-chromosomal inversions, but differ in their TE content with Lmb having undergone a massive TE expansion 5 million years ago corresponding to the speciation date (Grandaubert *et al*., 2014). Invasion of TEs in the genome of Lmb has shaped its genome, with alternating TE-rich regions and gene-rich regions. In contrast, the Lml genome shows a homogeneous TE distribution along the chromosomes (Grandaubert *et al*., 2014).

The distinct genome organisation shown by Lmb and Lml thus provides us with a model of choice for comparing epigenomic organization in two closely related phytopathogenic fungi, to determine the genomic location of pathogenicity/effector genes in relation to the chromatin landscape and the influence of chromatin structure on gene expression. Comparative epigenomic analyses, at the intra- or inter-species levels, are very sparse and this remains an underexplored field of study, at least in fungi (Zhang *et al*., 2018; Feurtey *et al*., 2019). We present here the first comparative epigenomic analysis of both euchromatin and heterochromatin marks in two closely related phytopathogenic fungi. We first performed ChIP-seq and RNA-seq during axenic culture to compare the distribution of three histone modifications, H3K4me2, H3K9me3 and H3K27me3, and then investigated whether chromatin organization is conserved between Lmb and Lml by assessing whether orthologous genes, and pathogenicity-related genes, are located in similar chromatin domains in each species. Lastly, we assessed the influence of the chromatin landscape on gene expression during axenic growth, focusing mostly on pathogenicity related genes that may be species-specific.

## Materials and Methods

### Fungal isolates

The isolate v23.1.3 of *L. maculans* ‘brassicae’ and the isolate IBCN84 of *L. maculans* ‘lepidii’ were used throughout the analyses (Rouxel *et al.*, 2011; Grandaubert *et al.*, 2014). Fungal cultures were maintained as described previously (Ansan-Melayah *et al*., 1995). For chromatin immunoprecipitation and transcriptomic analyses performed during *in vitro* growth, mycelium of Lmb and Lml were grown on V8 agar medium at 25°C for seven days. Then 10 plugs of mycelium were inoculated into 100 ml of Fries liquid medium in Roux bottle. Mycelia were harvested after growing for 7 days at 25°C, filtered and washed thoroughly with distilled water, and immediately placed in liquid nitrogen until further used. ChIP experiments were performed on fresh material.

### Chromatin immunoprecipitation and high-throughput sequencing

ChIP was performed from freshly-harvested mycelium grown in Fries liquid culture, as described in Soyer *et al*. (2015b), with minor modifications. ChIP was performed on native material (without crosslinking) using antibodies targeting histone modifications H3K4me2 (Merck ref. 07-030), H3K9me3 (Active Motif, Carlsbad, CA, USA; ref. 39161), or H3K27me3 (Active Motif, Carlsbad, CA, USA; ref. 39155). Three different ChIPs (i.e. three biological replicates) were performed for each of the histone modifications. Libraries were prepared from all biological replicates, individually, according to the Illumina TruSeq protocol “Ultra Low Input DNA library”. Libraries were sequenced with an Illumina HiSeq 2000 genome analyzer at the Max Planck Genome centre Cologne, Germany (https://mpgc.mpipz.mpg.de/home/). Sequencing data are available under the GEO accession number GSE150127.

### Analysis of ChIP-seq data and identification of significantly enriched domains

Analysis of ChIP-seq datasets was performed as described in Schotanus *et al*. (2015). Quality of Illumina reads was analysed using FastQC (https://www.bioinformatics.babraham.ac.uk/projects/fastqc/). Based on results of this analysis, ten bp were trimmed from the 5’ end. Processed reads were mapped on the reference genome of Lmb (Dutreux *et al*., 2018) or Lml (Grandaubert *et al*., 2014) using Bowtie2 (Langmead and Salzberg, 2012) with default parameters (Table S1). Peak calling analysis was performed on each ChIP sequencing dataset, to identify significantly enriched domains for either H3K4me2, H3K9me3 or H3K27me3, using RSEG (Song and Smith, 2011). A domain was considered if identified in at least two out of the three biological replicates. The Integrative Genome Viewer (Thorvaldsdóttir *et al.*, 2013) was used to visualize location of each domain along genomes of Lmb and Lml according to other genome features (gene annotation, TE-annotation, GC-content). Coverage of the histone modifications was assessed in 10 kb non-overlapping sliding windows along the supercontigs and correlation analyses, using Kendall’s 𝒯 correlation coefficient, were performed between biological replicates to check for reproducibility (Tables S2, S3). In order to assess significant enrichment of H3K4me2, H3K9me3 or H3K27me3 in certain categories of genes (such as proteinaceous or metabolic effector-encoding genes), a Chi^2^ test was applied to compare the expected proportion of a given category of genes across the entire genome to the observed distribution of the gene category in H3K4me2-, H3K9me3- or H3K27me3-domains (Soyer *et al*., 2019). Enrichment was considered significant with a *P value* < 0.01; analyses were done using R, version 3.0.2 (www.r-project.org).

### RNA extraction, RNA-sequencing and expression analysis

Total RNA was extracted from mycelium grown for one week in Fries liquid medium as previously described (Fudal *et al*., 2007). The NEBNext Ultra Directional RNA Library Prep Kit for Illumina (cat. # E7420L New England BioLabs) was used to prepare RNA-seq libraries and sequencing was performed on a HiSeq 4000 Illumina genome analyser using 50 paired-end reads. Raw reads were then pre-processed with Trimmomatic (Bolger *et al*., 2014) to remove short reads (<30 bp) and eliminate sequencing adaptors. Cleaned reads were then mapped against each genome using STAR with default parameters (Dobin and Gingeras, 2015; Table S1). Gene expression was evaluated using the rpkm_count function of EdgeR (Robinson *et al*., 2010). Genes with RPKM ≥ 2 were considered expressed. Sequencing data are available under the GEO accession number GSE150127.

### Identification of orthologues and annotation of genes encoding proteinaceous effectors in Lmb and Lml

The genome of Lmb encodes 13,047 proteins and that of Lml 11,272 proteins (Grandaubert *et al.*, 2014; Dutreux *et al.*, 2018). Orthologous proteins between Lmb and Lml were identified using OrthoFinder with default parameters (Emms and Kelly, 2015).

Based on the new assembly and annotation available for the genome of Lmb (Dutreux *et al*., 2018), an updated repertoire of putative proteinaceous effector genes (encoding Small Secreted Proteins, SSP) has been predicted. Therefore, to compare effector repertoires of Lmb and Lml, the same pipeline for the prediction of putative effector genes was applied to both species. Signal peptide and subcellular localization were predicted by SignalP version 4.1 (Petersen *et al*., 2011) and TargetP version 1.1 (Almagro Armenteros *et al*., 2019) respectively. Transmembrane domains were predicted by TMHMM version 2.0. The predicted secretome contained all proteins with no more than one transmembrane domain and either a predicted signal peptide or a predicted extracellular localization. The final effector repertoires were created by applying a size cut-off of 300 amino acids on the predicted secretome. In parallel, EffectorP version 1.0 (Sperschneider *et al.*, 2016) was used on the predicted secretome and, for Lml, four proteins were predicted as effectors with a size higher than 300 amino acids. These four proteins were added to the SSP set. This predicted a repertoire of 1,080 and 892 SSP-encoding genes for Lmb and for Lml, respectively.

### Analysis of GO enrichment

Gene Ontology (GO) annotations of the Lmb and Lml genes were retrieved from Dutreux *et al*. (2018) and Grandaubert *et al*. (2014) respectively. GO term enrichment analysis of the H3K4me2-, H3K9me3- and H3K27me3-associated genes was performed with the plug-in Biological networks Gene ontology (BinGo; v3.0.3) of the cytoscape software (Shannon *et al*., 2003). List of genes submitted to BINGO were considered as significantly enriched for a given GO term with an associated False Discovery Rate (FDR) ≤ 0.01 for the biological processes.

A Chi^2^ test, or a Fisher’s exact test for small sized-population, were applied to identify significant enrichment of H3K4me2-, H3K9me3- or H3K27me3-domains for certain categories of genes: the expected proportion of a given category of genes across the entire genomes of Lmb or Lml was compared to the observed distribution of the gene category in the H3K4me2-, H3K9me3- or H3K27me3-domains. Enrichment was considered significant with a *P value* < 0.01. All analyses were done using R, version 3.0.2 (www.r-project.org).

## Results

### A different genome organisation but similar epigenomic properties in both genomes

To assess the genome-wide distribution of histone marks in Lmb and Lml during axenic growth, we performed ChIP-seq experiments using antibodies against histone modifications H3K4me2, H3K9me3 and H3K27me3 (Table S1). Mapping of the ChIP-seq data was followed by identification of significantly enriched domains, for any of the histone modifications targeted, and reproducibility of the biological replicates was assessed (Tables S2, S3). Based on the genomic coordinates of the H3K4me2-, H3K9me3- and H3K27me3-domains, the number of bases associated with any of the three histone modifications was evaluated, for each genome, in 10 kb sliding-windows. In the genomes of Lmb and Lml, the proportion of H3K4me2 and H3K27me3 was similar (H3K4me2: 39% in Lmb vs. 32% in Lml; H3K27me3: 19% in Lmb vs. 13% in Lml) while the proportion of H3K9me3 was strikingly different (33% and 4% for Lmb and Lml, respectively) (Tables S4, S5). While the median size of the H3K4me2- and H3K27me3-domains was similar in both genomes, the Lmb genome displayed extremely large H3K9me3-domains, with a domain encompassing up to 230 kb, while the maximum size of the H3K9me3-domains in Lml was only 12 kb (Figures 1; 2).

**Figure 1.**
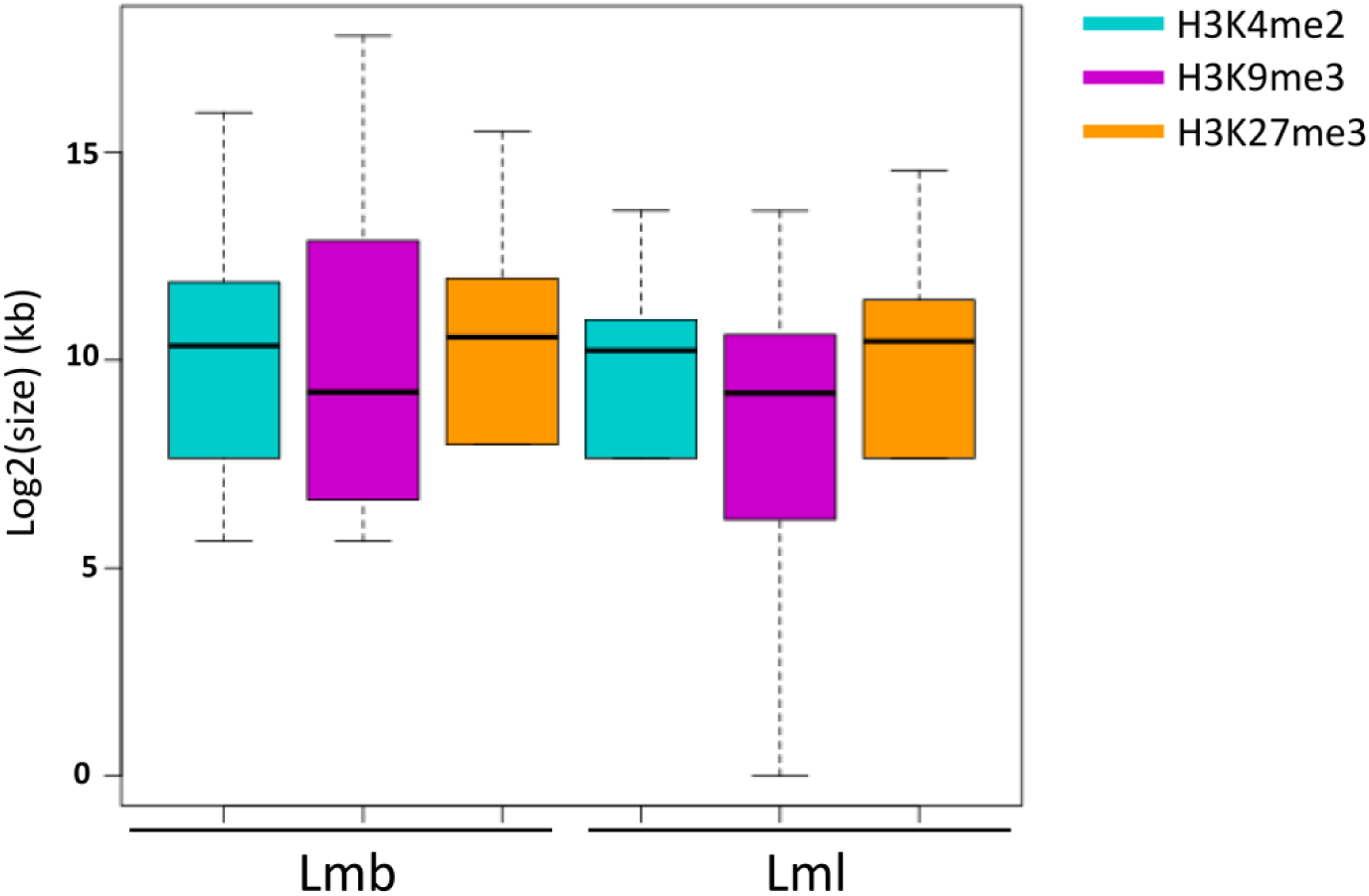
Size of the H3K4me2-, H3K9me3-, H3K27me3-domains in the genomes of *Leptosphaeria maculans* ‘brassicae’ (Lmb) and *L. maculans* ‘lepidii’ (Lml). Log2 of the size of the domains was estimated based on the coordinates of the location of the domains, identified using RSEG (Song and Smith, 2011). Blue: H3K4me2; Purple: H3K9me3; Orange: H3K27me3.

**Figure 2.**
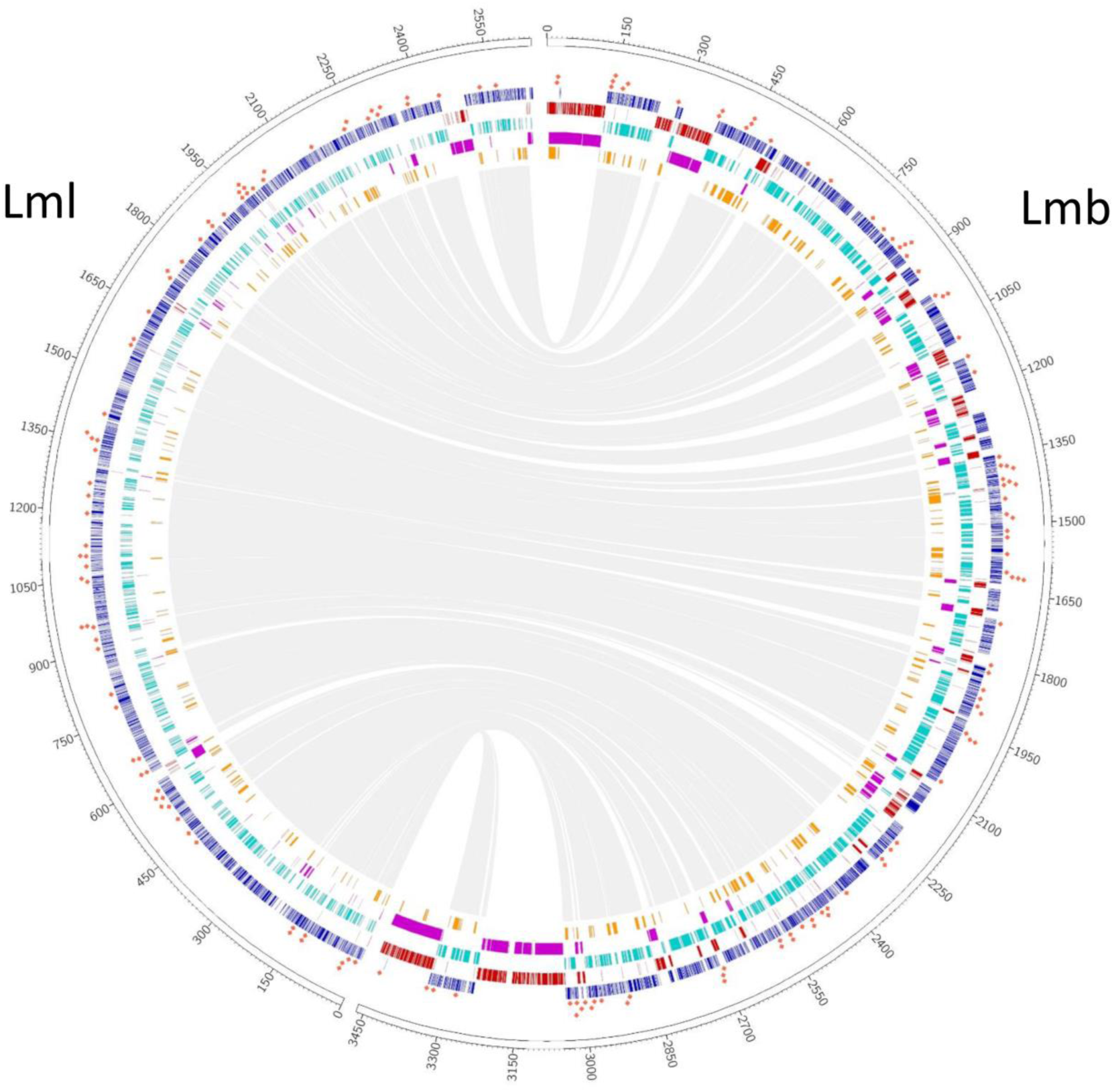
Genome of *Leptosphaeria maculans* ‘brassicae’ (Lmb) harbors large TE-rich, H3K9me3-domains compared to *Leptosphaeria maculans* ‘lepidii’ (Lml). Example of SuperContig 2 of Lmb and Scaffold 1 of Lml. ChIP-seq was performed with antibodies targeting H3K4me2 (cyan), H3K9me3 (purple) or H3K27me3 (orange); rectangles indicate location of significantly enriched domains identified using RSEG (Song and Smith, 2011). Blue: location of CDS; red: location of transposable elements. Genes encoding proteinaceous effectors are indicated with a red square.

In the genomes of Lmb and Lml, H3K4me2- and H3K9me3-domains were mutually exclusive (Kendall’s *𝒯*: −0.57 and −0.16, *P* < 2.2.10^−16^ for Lmb and Lml respectively; Figure 2; Figure 3; Tables S6 and S7). We identified a positive correlation between the location of TEs and H3K9me3-domains (Kendall’s *𝒯*: 0.87 and 0.68, *P* < 2.2.10^−16^ for Lmb and Lml respectively; Figure 2; Figure 3; Figure 4A, 4C; Table S6 and S7) and between the location of coding sequences (CDS) and H3K4me2 (Kendall’s *𝒯*: 0.55 and 0.25, *P* < 2.2.10^−16^ for Lmb and Lml respectively; Figure 2; Figure 3; Figure 4A, 4C; Table S6 and S7). In the genome of Lmb, a weak correlation was demonstrated between CDS and H3K27me3-domains (Kendall’s *𝒯*: 0.12, *P* < 2.2.10^−16^; Figure 3; Tables S6, S7). For Lmb and Lml, the coverage of H3K9me3 and H3K27me3 was not homogeneous across all SCs because some are >2-fold more highly enriched with these histone modifications compared to others. An extreme example of this situation was identified on the dispensable chromosome of Lmb (Leclair *et al*., 1996; Rouxel *et al*., 2011), which is extremely enriched in TEs compared to the rest of the genome (35% of TEs in the core genome and 93% of TEs in the dispensable chromosome). Consistently, a strong enrichment in H3K9me3 was observed in the dispensable chromosome (32% in the core genome and 90% in the dispensable chromosome), with only a few H3K4me2- and H3K27me3-domains (Figure 4A and 4B). In Lmb, we found that many sub-telomeric regions showed overlaps between H3K9me3 and H3K27me3, whereas these marks did not overlap on the chromosome arms, even within large TE-rich regions (Figure 2). The assembly quality of the Lml genome was not sufficient to allow us to evaluate whether this was a common feature between the two species.

**Figure 3.**
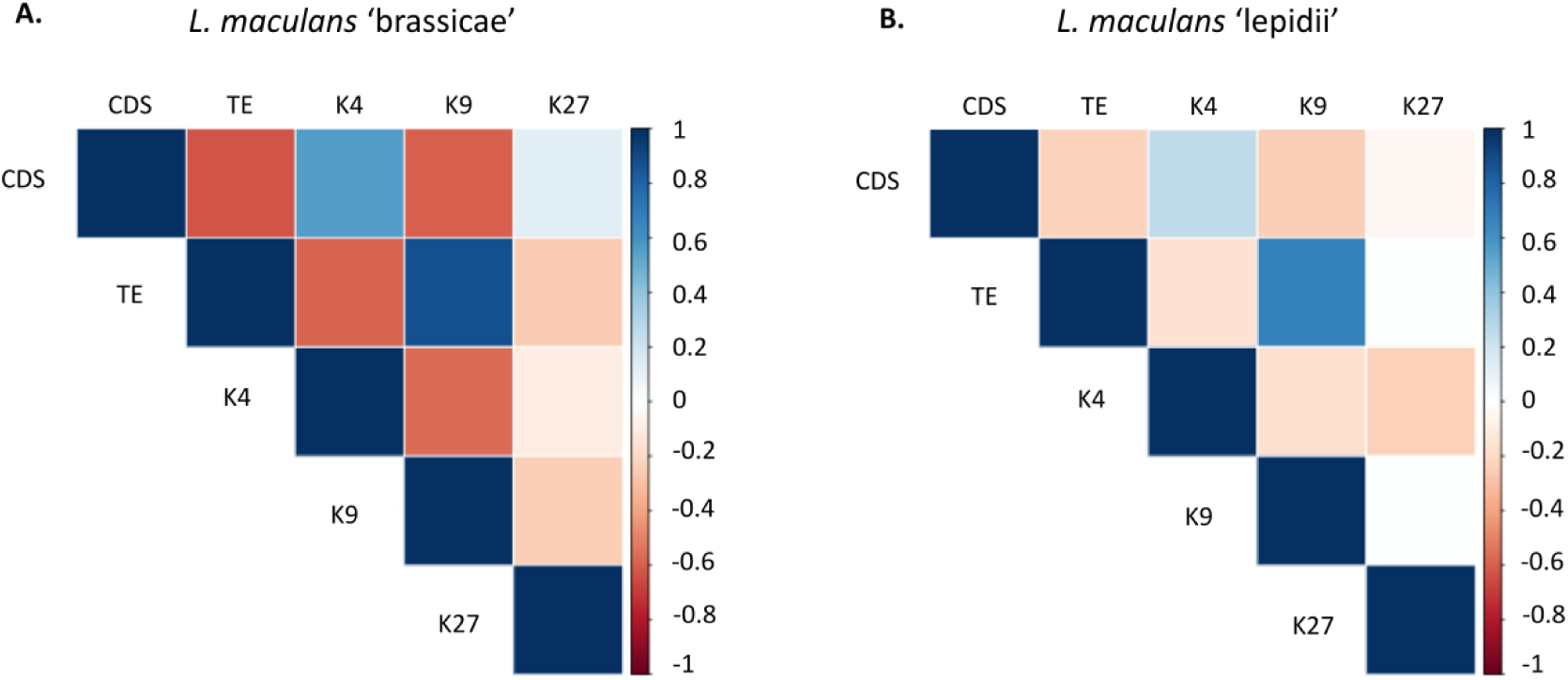
Correlation analysis of the location of genes, transposable elements and domains enriched for H3K4me2, H3K9me3 and H3K27me3 in the genomes of A. *Leptosphaeria maculans* ‘brassicae’; B. *Leptosphaeria maculans* ‘lepidii’. Regions significantly enriched for any of the three histone modifications analysed were identified through ChIP-seq performed during axenic culture; correlation analyses were performed using a Kendall’s *𝒯* (see Materials and Methods). K4: H3K4me2; K9: H3K9me3; K27: H3K27me3; CDS: Coding sequences; TE: Transposable Elements.

**Figure 4.**
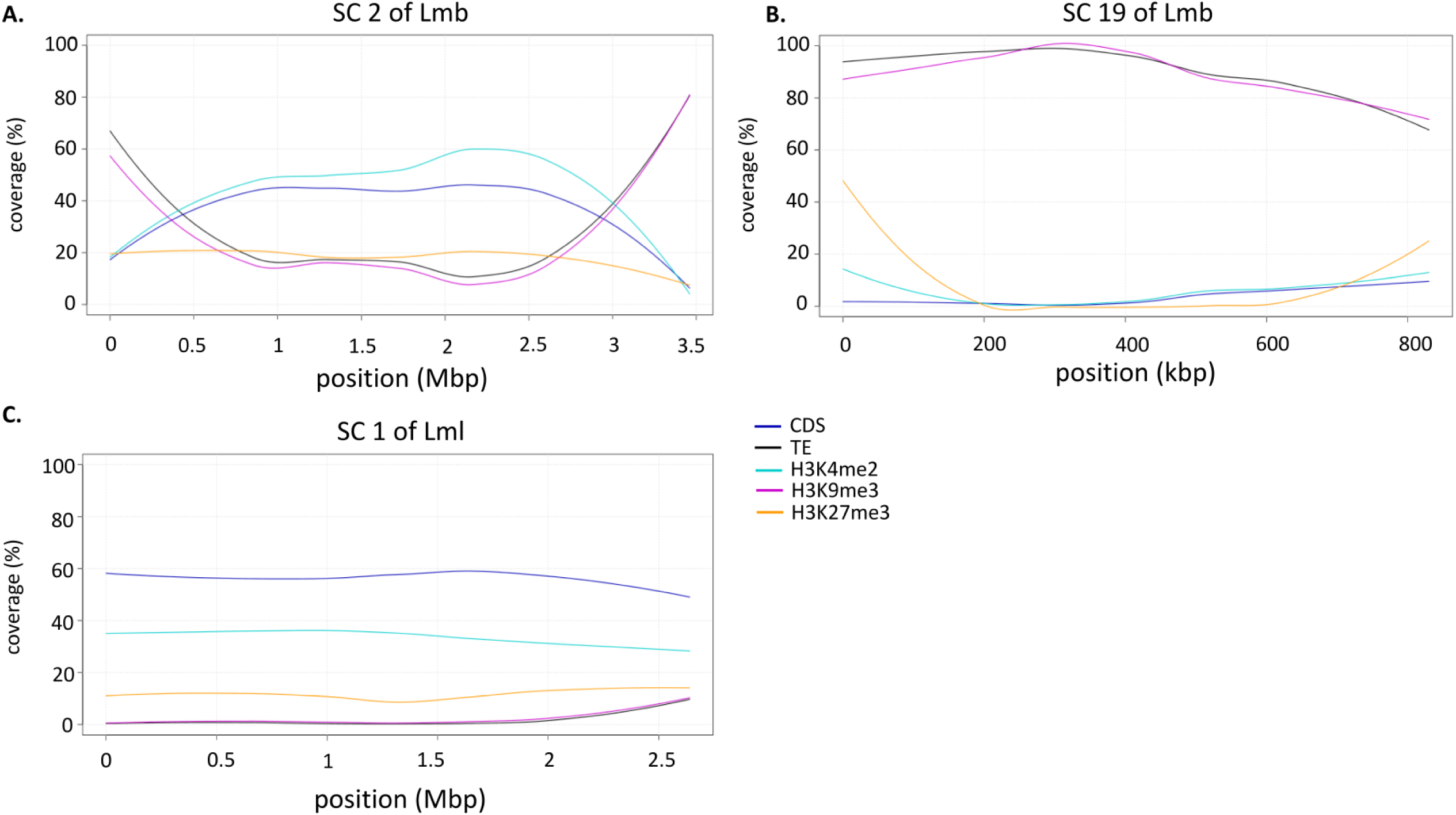
Coverage of genes, transposable elements, H3K4me2-, H3K9me3- and H3K27me3-domains in the genomes of *L. maculans* ‘brassicae’ (Lmb) and *L. maculans* ‘lepidii’ (Lml). A. SuperContig 2 of Lmb; B. SC 19, i.e. dispensable chromosome, of Lmb; C. SC 1 of Lml. SC2 of Lmb and SC1 of Lml are syntenic (Grandaubert *et al*., 2014). Coverage of the genome features and histone modification domains, as identified in axenic cultures, were analysed in 10 kb sliding windows in the genomes of Lmb and Lml. CDS, Coding Sequences; TE, Transposable Elements; SC, Super Contig.

### H3K4me2-domains associate with genes involved in primary metabolism while H3K27me3-domains shelter genes involved in niche adaptation and cell wall degradation

Genes associated with H3K4me2-, H3K9me3- or H3K27me3-domains were identified in the genomes of Lmb and Lml during axenic growth. Overall, the number of genes associated with any of the histone post-translational modifications was similar in both species, with more than 50% of the predicted genes associated with H3K4me2, ∼14% of the genes associated with H3K27me3 and only a few genes associated with H3K9me3 (104 and 70 respectively in Lmb and Lml; Figure 5). In Lmb, there was a higher number of genes associated with both H3K4me2 and H3K27me3 than in Lml (2,044 and 361 for Lmb and Lml, respectively) (Figure 5). In both genomes, a GO annotation could be assigned to 30-40% of the predicted genes (5,076 and 3,725 genes for Lmb and Lml, respectively; Grandaubert *et al*., 2014; Dutreux *et al*., 2018). Euchromatin was enriched with genes with a GO annotation (X^2^ test, *P* < 2.2.10^−16^) because more than 40% of the genes associated with H3K4me2 had a GO annotation (3,276 and 2,581 genes for Lmb and Lml respectively). Between 20 and 28% of the genes associated with H3K27me3 (572 and 288 genes for Lmb and Lml, respectively) had a GO annotation. However, none of the genes located in H3K9me3-domains, H3K9/K27me3-domains or H3K4me2/H3K27me3-domains had a GO annotation. In other words, genes located in heterochromatin were enriched with genes lacking any functional annotation (X^2^ test, *P* < 2.2.10^−16^). For both species, genes associated with H3K4me2 displayed a wide variety of annotated functions corresponding to primary metabolism and basic cellular functions, such as translation (GO:0006412; 206 genes) or cellular protein metabolic process (Tables S8, S9). For both species, only a few GO terms were identified among genes associated with H3K27me3. Nevertheless, in Lmb these genes were significantly enriched in GO terms associated with carbohydrate metabolic process (GO:0005975; 82 genes), oxydo-reduction process (GO:0055114; 149 genes) and transmembrane transport (GO:0055085; 104 genes) (*P* ≤ 0.01) (Table S10). As for Lml, probably due to the lower number of genes associated to H3K27me3, only a few GO annotation enrichments were detected. Only one GO enrichment was found in common with Lmb, namely carbohydrate metabolic process (GO:0005975; 41 genes). Other enrichments corresponded to a few genes classified as response to chemical (GO: 42221; 20 genes), response to nitrogen compound (GO:1901698; 10 genes), response to organophosphorus (GO: 46683, 10 genes) and others (Table S11). Although most GO enrichments found among sets of genes associated with H3K27me3 in Lmb and Lml did not overlap, both types of enrichments suggest that H3K27me3-associated genes might be involved in stress response mechanisms, but may also be involved in feeding or cell wall degradation processes during plant infection.

**Figure 5.**
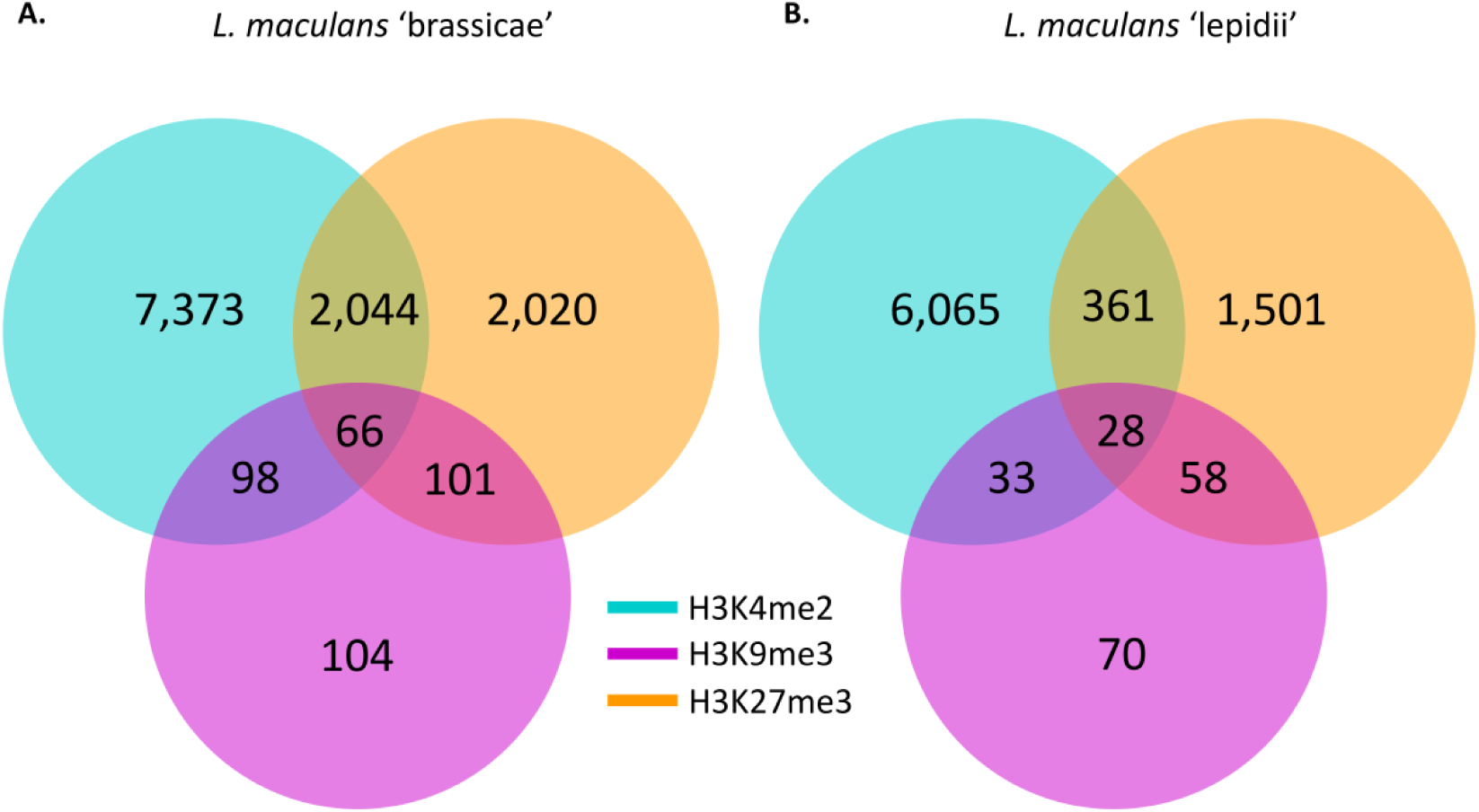
Number of genes associated with H3K4me2, H3K9me3 or H3K27me3 during axenic growth in A. *Leptosphaeria maculans* ‘brassicae’ (Lmb) and B. *L. maculans* ‘lepidii’ (Lml). Locations of histone modifications in the genomes of Lmb and Lml were identified using RSEG (Song and Smith, 2011). Blue: H3K4me2; Purple: H3K9me3; Orange: H3K27me3. Genes were considered as associated with any of the histone modifications when at least one pb of the gene was found within the borders of the domain.

### H3K4me2-domains are associated with expressed genes while H3K9me3- and H3K27me3-domains are associated with silent genes during axenic growth

We performed a genome-wide transcriptomic analysis of fungal cultures grown *in vitro*, and we correlated gene expression patterns with the distribution of histone modifications. In Lmb, 10,934 genes (83% of the predicted genes) and 7,735 genes in Lml (69% of the predicted genes) were expressed during vegetative growth. Considering the top 100 most expressed genes of Lmb, 68 were associated with H3K4me2 while two were located within a H3K27me3-domain and none in a H3K9me3-domain (data not shown). In Lml, 71 were located within a H3K4me2-domain while five were located in a H3K27me3-domain (data not shown). These data were confirmed at a genome-wide scale because H3K4me2-domains were enriched with genes expressed during axenic culture (7,104 of the genes associated with H3K4me2 were expressed in Lmb and 5,338 in Lml; Figure 6; *P* < 2.2.10^−16^). On the contrary, H3K9me3- and H3K27me3-domains were enriched with genes that were silent during *in vitro* growth (Figure 6; *P* < 2.2.10^−16^). On the other hand, in both species, genes that were associated with both H3K4me2 and H3K27me3 were expressed during axenic growth (90% and 87% of the genes, respectively, in Lmb and Lml) and their level of expression was similar to that of genes located in H3K4me2-domains (Figure 6). In both species, this transcriptomic analysis confirmed that genes located in a H3K4me2-domain were more likely to be expressed while those located in H3K9me3- and H3K27me3-domains were more likely to be silent.

**Figure 6.**
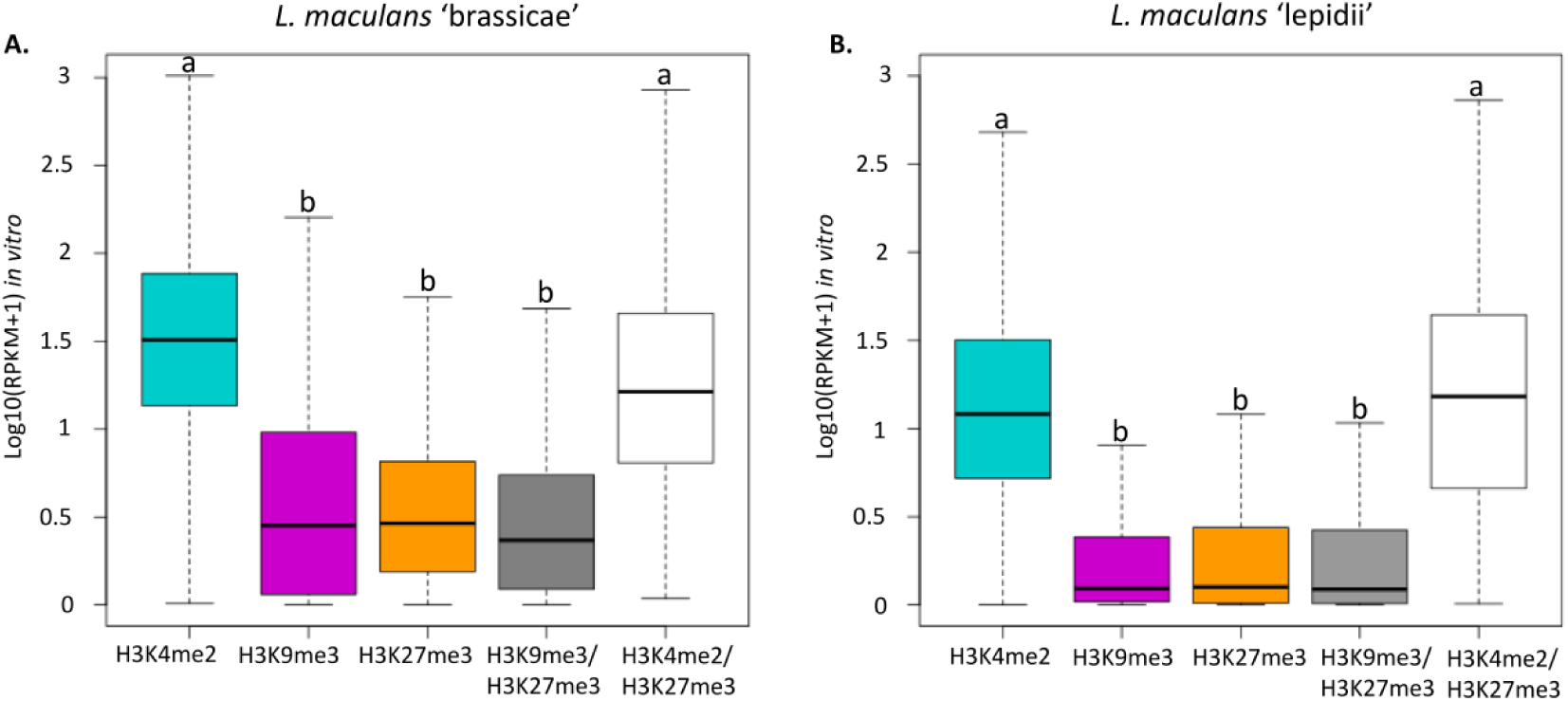
Genes associated with heterochromatin are less expressed than genes associated with euchromatin in the genomes of A. *L. maculans* ‘brassicae’ and B. *L. maculans* ‘lepidii’. Location of histone modifications in the genome of Lmb and Lml were identified using RSEG (Song and Smith, 2011), during axenic culture. RNA-seq was performed from Lmb or Lml grown one week in FRIES media. Blue: H3K4me2; Purple: H3K9me3; Orange: H3K27me3.

### Heterochromatin domains are enriched with species-specific genes

A total of 7,393 genes were conserved between both species. H3K4me2-domains were enriched with genes conserved between these two species, for example 4,892 Lmb genes located in euchromatin regions during growth *in vitro* were conserved (X^2^ test, *P* < 2.2.10^−16^) and the vast majority of these (4,298, i.e. 88%) were expressed during axenic growth in both Lmb and Lml. Most of the conserved genes associated with H3K4me2 were involved in primary metabolism as mentioned above. Genes associated with both H3K4me2 and H3K27me3 were also enriched with conserved genes (1,318 and 262 genes, respectively, in Lmb and Lml) representing more than 65% of the genes in these domains (X^2^ test, *P* < 5.2.10^−3^). Two genes involved in heterochromatin assembly and maintenance were analysed previously through gene silencing (*LmHP1* and *LmKMT1*/*DIM5*; Soyer *et al*., 2014). *LmHP1* is conserved in Lml and located in euchromatin in both species. *LmKMT1* is also conserved in both species, being located in a H3K4me2/H3K27me3 domain in Lmb and within a H3K4me2/H3K9me3 domain in the case of Lml, during axenic growth. In contrast, H3K27me3- and H3K9me3-domains were significantly enriched with species-specific genes (a total of 865 genes and 661 genes, respectively, in Lmb and Lml; X^2^ test, *P* < 2.2.10^−16^). Among genes located in heterochromatin in both species (either H3K9me3-, H3K27me3- or H3K9/K27me3-domains), 445 were conserved between Lmb and Lml; of which 83 were expressed and 182 were repressed during axenic growth in both species. No GO enrichment was found for these genes and predicted functions were sparse, with less than 45% of them having a functional annotation. Strikingly, the main enrichment was in genes encoding putative effectors (16% of the genes).

### Heterochromatin domains are enriched with proteinaceous and metabolic effector genes

We then focused on candidate proteinaceous or metabolic effectors and wondered whether they were conserved and showed a distinct pattern of histone modifications. In the genome of Lmb, 2,478 genes (i.e. 12% of the total genes) were associated with TE-rich regions (i.e. located within 2 kb distance of a TE sequence), of which 289 genes encoded putative proteinaceous effectors. Hence, although a new prediction of the effector repertoire was performed here, based on the new assembly of the Lmb genome, TE-rich regions were significantly enriched with proteinaceous effector genes, as was already shown by Rouxel *et al*. (2011; X^2^ test, *P*= 9.6.10^−16^). Similarly, both H3K9me3- and H3K27me3-associated genes were significantly enriched with putative proteinaceous effectors, as they represented, respectively, 36% and 14% of the genes associated with these histone modifications *in vitro* (X^2^ test, *P*< 2.2.10^−16^; Table 1). Likewise, in Lml, TE-rich regions, H3K9me3- and H3K27me3-domains were significantly enriched with putative proteinaceous effector genes while H3K4me2-domains were significantly depleted in such genes compared to the rest of the genome (Table 1). Among the 1,080 putative proteinaceous effector genes predicted in Lmb (8.2% of the total predicted genes) and the 892 putative effector genes of Lml (7.9% of the total predicted genes), 274 were conserved between both species. Hence, more than two-thirds of the effector repertoire is species-specific (X^2^ test, *P* < 2.2.10^−16^), confirming our previous findings (Grandaubert *et al.*, 2014). Overall, orthologous effector-encoding genes were associated with the same types of chromatin domains in Lmb and Lml. For example, 98% of the Lmb effector genes located in a euchromatin environment were also associated with H3K4me2 in Lml (82% of Lml effector genes located in euchromatin regions were also associated with H3K4me2 in Lmb). H3K27me3- and H3K9me3-domains were enriched with species-specific effector genes in both species (Table 1). As a striking example of the non-random location of effector genes in the two genomes, and the enrichment of heterochromatin regions with species-specific effectors, all nine currently known avirulence genes of Lmb were located in H3K9me3-domains *in vitro*, consistent with the fact that these genes are located in TE-rich genomic compartments; none of the nine was conserved in the genome of Lml. Twenty-eight other proteinaceous effector-encoding genes were located in TE-rich, H3K9me3-domains in Lmb, and only one of them had a putative ortholog in Lml. The 274 orthologous genes encoding effectors were then investigated for their distribution in euchromatic/heterochromatic regions during vegetative growth and for conservation of their location between orthologs. There was no obvious bias in the distribution of chromatin marks among the 274 genes, which is comparable to the overall distribution of marks among all genes of the effector repertoires (data not shown). At the individual gene level, 86 of the 274 orthologs were located in a similar chromatin domain in Lmb and Lml, including 83 genes associated with H3K4me2 and 47 genes associated with H3K27me3 in both genomes. In contrast to avirulence genes, ‘late’ putative effector-genes of Lmb (i.e. expressed during stem colonization) were located outside of TE-rich regions (Gervais *et al*., 2017). Three of the 11 experimentally validated ‘late’ effector genes were located in a H3K4me2-domain while others were located in a H3K9me3- (two cases), a H3K27me3- (three cases), a H3K9/K27me3- (one case) or a H3K4me2/K27me3-domain (two cases), suggesting that ‘late’ effector genes were also enriched with heterochromatin domains during growth *in vitro*. Interestingly, and contrary to the trend observed for Lmb avirulence effector genes (and other effector candidates associated with TE-rich regions), only two of the 11 ‘late’ putative effector genes, either located in H3K9me3- or H3K27me3-domains, were not conserved in Lml. Taken together, these data show that, independently of the species or the stage at which they are expressed during infection, proteinaceous effector genes are enriched in H3K9me3- or H3K27me3-domains during axenic growth.

**Table 1.**
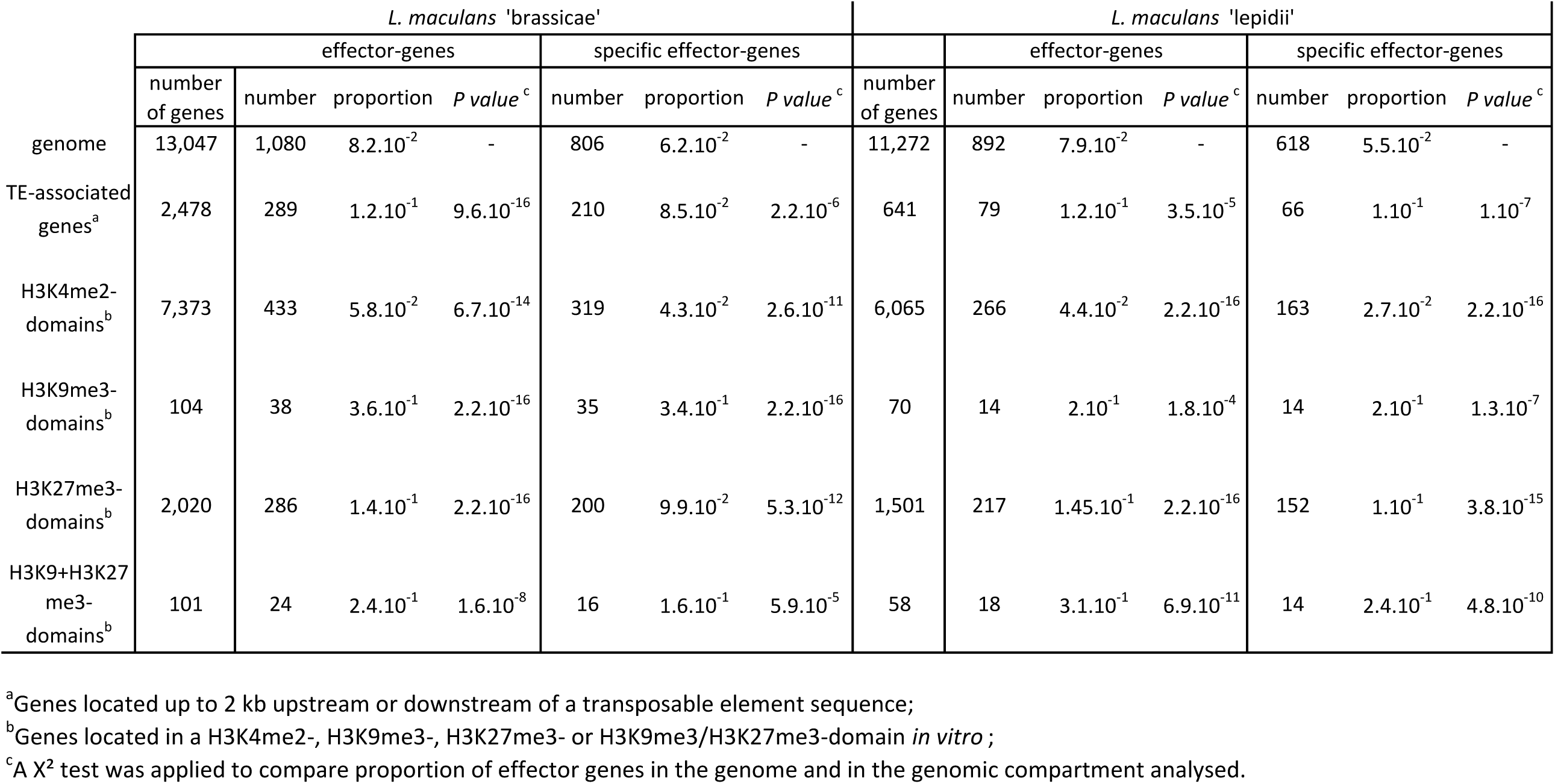
Number of proteinaceous effector genes located in different genomic compartments in *L. maculans* ‘brassicae’ and *L. maculans* ‘lepidii’.

The genomes of Lmb and Lml contain secondary metabolite gene clusters including key genes encoding PKS (Polyketide Synthases) and NRPS (Non-Ribosomal Peptide Synthetases). Twenty-seven such genes were predicted in Lmb among which 24 were conserved in Lml, but they were overall absent from other closely related species (Rouxel *et al*., 2011; Grandaubert *et al*., 2014; Table S12). Of these, only three have been experimentally demonstrated to be involved in Lmb pathogenicity, namely the PKS responsible for synthesis of abscisic acid (ABA), only expressed during cotyledon infection (Darma *et al*., 2019), the PKS responsible for producing phomenoic acid (Elliott *et al*., 2013), and the NRPS responsible for synthetising sirodesmin, a toxin produced during stem infection (Gardiner *et al*., 2004; Elliott *et al*., 2007). The ABA PKS and all seven genes of the cluster, which are intermingled with three TE-rich regions, were entirely absent from the genome of Lml, while the other two were conserved. In Lmb, 81% of the PKS/NRPS-encoding genes were associated to H3K27me3- or H3K4me2/H3K27me3-domains (22 PKS/NRPS; Fisher’s exact test, *P*= 1.3.10^−7^). Likewise in the Lml genome, H3K27me3-domains were enriched with PKS/NRPS-encoding genes (12 PKS/NRPS; Fisher’s exact test, *P*= 1.7.10^−4^), and all of them had orthologs associated with similar marks in Lmb. The similar chromatin context also resulted in similar regulation in most of the cases, with 19 of the orthologs being similarly expressed during *in vitro* growth (17 expressed and two repressed).

## Discussion

The sister species *L. maculans* ‘brassicae’ and *L. maculans* ‘lepidii’ exhibit marked differences in their genome organisation. Notably, Lmb has large TE-rich domains structuring the genome into alternating gene-rich and TE-rich regions (Rouxel *et al*., 2011). In the genome of Lml, in contrast, TEs are evenly distributed across the genome and no compartmentalisation of the genome is evident in relation to TE location (Grandaubert *et al*., 2014). The massive invasion of the Lmb genome by TEs occurred ca. 5 MYA and was postulated to have been instrumental in the separation of the two species (Grandaubert *et al*., 2014). This invasion may also have contributed to the rise of Lmb as a successful pathogen of *B. napus* due to the specific localisation of a number of candidate effector genes in TE-rich regions of the genome (Rouxel *et al*., 2011; Grandaubert *et al*., 2014). Differences in genomic organization between these two closely-related species could impact the underlying epigenomic landscape and have important consequences for fungal biology and pathogenicity. This could provide distinct strategies to regulate the expression of genes involved in stress response or pathogenicity, as it was shown in Lmb that histone modification H3K9me3 is involved in the repression of avirulence genes during axenic growth (Soyer *et al*., 2014). To investigate this question, we here compared the genome-wide location of three different histone modifications that are typically associated with euchromatin or heterochromatin in these two closely-related phytopathogenic fungi. We found that differences in the epigenomic landscape of Lmb and Lml are in accordance with their genome organization. In Lmb, very large H3K9me3-domains are present, spanning large TE-rich regions, while such extremely large H3K9me3-domains are not observed in the Lml genome. Our findings corroborate previous epigenetic analyses of a few genes performed *in vitro*, pointing out that avirulence genes are located in heterochromatin, but also demonstrate that putative effector genes, independently of their expression pattern or their location in TE-rich or gene-rich regions, are enriched with heterochromatin during axenic culture.

Although Lmb and Lml have distinct genome organisations, they share a common distribution of histone modifications throughout their genome during axenic culture. Gene-rich regions are enriched with H3K4me2 and H3K27me3, while TE-rich regions are associated with H3K9me3. The proportion of H3K9me3 in a genome often reflects the TE content, as was described in *Z. tritici* (Schotanus *et al*., 2015). In Lmb, having more than 30% of TE, 33% of the genome is associated with H3K9me3 while the genome of Lml, having a low TE content, shows a low enrichment in H3K9me3 (4% of H3K9me3). Domains enriched with H3K4me2 and domains enriched with H3K9me3 are mutually exclusive in the genomes of Lmb and Lml, as was shown in *Neurospora crassa, Fusarium fujikuroi*, and *Z. tritici* (Smith *et al*., 2008; Jamieson *et al*., 2013; Wiemann *et al*., 2013; Schotanus *et al*., 2015). H3K27me3 has been detected in most filamentous fungi investigated so far, except in *Mucor, Rhizopus* or Aspergilli such as *Aspergillus nidulans* (Gacek-Matthews *et al*., 2016; reviewed in Freitag, 2017). In sub-telomeric regions of Lmb, H3K9me3 and H3K27me3-domains overlap over repetitive sequences, which is also the case in *N. crassa* or *Z. tritici* (Smith *et al*., 2008; Jamieson *et al*., 2013; 2017; Schotanus *et al*., 2015). In the genome of Lmb, TE-rich regions are enriched with H3K9me3 but, except for the sub-telomeric regions, no enrichment in H3K27me3 was associated with TEs on chromosomal arms. This contrasts with the organization of TE-rich regions in *Z. tritici*, which are enriched with both heterochromatin modifications (Schotanus *et al*., 2015). The situation in Lmb is also different from that of *Fusarium oxysporum* in which most TE sequences are embedded in H3K27me3-domains (Fokkens *et al*., 2018). We confirmed that, as in most Eukaryotes, H3K4me2-domains are associated with expressed genes while H3K9me3- and H3K27me3-domains are associated with silent genes. In Lmb, although the locations of H3K4me2- and H3K27me3-domains do not overlap at a genome-wide scale, more than 2,000 genes were found to be associated with both histone methylations. This is strikingly different to Lml, where only 300 genes were associated with both histone modifications, and *Z. tritici*, where 400 genes are embedded in such domains (Schotanus *et al*., 2015; Soyer *et al*., 2019). Bivalent domains are defined as chromatin regions associated with both repressive and permissive histone modifications (Bernstein *et al*., 2006). The large number of genes associated with H3K27me3 and H3K4me2 during axenic culture in Lmb might indicate a biological specificity of this species and the existence of bivalent domains, because these two modifications have antagonistic effects on gene expression (Harikumar and Meshorer, 2015). Altogether, our findings show that, despite differences in genomic organization between Lmb and Lml (and the other fungi in which epigenomic analyses were so far performed), the epigenomic landscape is overall conserved.

While H3K9me3 and H3K27me3 are signatures of heterochromatin, H3K9me3 is considered to be typical of constitutive heterochromatin, being associated with repeats and involved in genome stability, whereas H3K27me3 is considered to be associated with facultative heterochromatin that is easily reversed to a euchromatin state under certain abiotic and biotic stress conditions (Grewal and Jia, 2007). However, dogmas regarding conventional definitions of facultative or constitutive heterochromatin seem to be challenged in fungi, or at least in the few plant pathogenic fungi in which epigenomic analyses have been performed. While the co-localization of H3K9me3 and TEs in the Lmb and Lml genomes is consistent with a constitutive heterochromatin state, the fact that H3K9me3-domains encompass genes that are expressed during interaction with oilseed rape suggests that these regions may correspond to a category of facultative heterochromatin in this species. Genes associated with H3K9me3 are almost all located in the middle of repeated elements, in sub-telomeric areas, or very close to the edge of regions enriched with repeated elements. Moreover, some of these genes, including the nine cloned avirulence genes of Lmb which are located within AT-rich isochores (e.g., *AvrLm1* or *AvLm6*; Gout *et al*., 2006; Fudal *et al*., 2007) including subtelomeric regions (e.g. *AvrLm3* or *AvrLm10*; Plissonneau *et al*., 2016; Petit-Houdenot *et al*., 2019), are highly transcribed upon host infection (Rouxel *et al*., 2011). This finding, together with other studies, questions the “constitutive” nature of heterochromatin associated with H3K9me3. In contrast, the location of H3K27me3 in the genomes of Lmb and Lml supports its association with facultative heterochromatin. Indeed, we found H3K27me3 associated with coding sequences and enriched with genes encoding proteinaceous and metabolic effectors or proteins involved in stress responses. Most of these genes are silenced during vegetative growth but induced during plant infection. In fungi, recent analyses also highlighted a role for H3K27me3 in genome organization and stability. For instance, the dispensable chromosomes of *Z. tritici* are twice as rich in TEs as the core chromosomes, and whereas there is no significant enrichment in H3K9me3 on the dispensable chromosomes, they are entirely covered by H3K27me3 (Schotanus *et al*., 2015). In *Z. tritici*, the loss of H3K27me3 was found to increase the stability of some accessory chromosomes (Möller *et al*., 2019). In *Z. tritici* and *N. crassa*, H3K27me3 is relocated towards normal constitutive heterochromatin (i.e. H3K9me3-domains) under genotoxic stress, such as the loss of H3K9me3 after inactivation of *KMT1* (Basenko *et al.*, 2015; Möller *et al*., 2019). Altogether, these findings suggest that although H3K27me3 is an important regulator of gene expression involved in development or response to various stresses, it also plays a role in the maintenance of genome integrity. In Lmb and Lml, no particular association of H3K27me3 with TEs was identified and the dispensable chromosome of Lmb is not enriched with H3K27me3. The only association of H3K27me3 with constitutive chromatin was at the chromosome ends of Lmb, where we found overlaps between H3K9me3 and H3K27me3. Inactivation of *KMT1* and *KMT6* would help investigate whether H3K27me3 could also be involved in genome stability and relocation of heterochromatin marks in Lmb and Lml.

The heterochromatin domains of Lmb and Lml are rich in species-specific genes, genes involved in stress responses and putative proteinaceous or metabolic effectors. This supports the view that chromatin remodelling mechanisms are an efficient way to rapidly modulate gene expression under stress conditions or during biotic interactions, although the dynamics of chromatin structure might not be the sole regulator of these complex biological processes (Soyer *et al*., 2015a; Tan and Oliver, 2017). This pattern is conserved in other plant-associated fungi, independently of their mode of interaction with their host (Ma *et al*., 2010; Rouxel *et al*., 2011; Chujo and Scott 2014; Schotanus *et al*., 2015; Faino *et al*., 2016; Vlaardingerbroek *et al*., 2016; Dallery *et al*., 2017; Krishnan *et al*., 2018; Soyer *et al*., 2019). Even outside of TE-rich regions, heterochromatin is often found enriched with pathogenicity-related genes, whether they encode proteinaceous effectors or are involved in the production of secondary metabolites (e.g. Connolly *et al*., 2013; Wiemann *et al*., 2013; Chujo and Scott, 2014; Schotanus *et al*., 2015; Fokkens *et al.*, 2018; Soyer *et al*., 2019). Importantly, sets of genes up-regulated during host infection are found associated with heterochromatin *in vitro* (Fokkens *et al*., 2018; Haueisen *et al*., 2018; Soyer *et al*., 2019). One of our initial postulates was that invasion of the *L. maculans* genome by TEs contributed to the rise of Lmb as a pathogen specialized on Brassicas with greater pathogenic abilities than the non-invaded sister species. While Lml has never been found to be able to infect *B. napus* under our experimental conditions or isolated from our experimental fields (M-H. Balesdent & T. Rouxel, unpublished data), previous work reported its ability to infect other crucifers (Petrie, 1969). Its isolation from ascocarps on stems of *Lepidium* sp. also suggests its infection strategy is similar to that of Lmb on *B. napus*. So far, all avirulence genes of Lmb, which can be recognized by the plant immune system to set up defence reactions, are located in TE-rich regions and associated with H3K9me3. In contrast, in both species we found conserved putative effector genes (either proteinaceous or metabolic) harbored in similar H3K27me3 heterochromatin environments. Moreover, ‘late’ effector genes are more conserved than avirulence genes between Lmb and Lml, and usually do not show presence/absence polymorphisms in Lmb (Gervais *et al.*, 2017). The contrasting heterochromatic environment (H3K9me3 vs. H3K27me3) for avirulence genes and ‘late’ effector genes reflects a very different adaptive behaviour because effector genes that are expressed early, and likely to be “recognised” by the plant surveillance machinery at the onset of penetration, are also those which are subject to accelerated evolution under selection (Rouxel *et al*., 2011). This suggests that there may be a selective advantage for the fungus to partition genes more likely to be recognised by the host plant within H3K9me3-domains. Nevertheless, the location of putative pathogenicity genes, including orthologues, in similar heterochromatin regions in the genomes of Lmb and Lml suggests that basic pathogenicity programs are independent of genome invasion by TEs, and points to the likewise importance of chromatin context on transcriptome shaping during infection. Taken together, these data support our previous hypothesis that the localization of effector genes in plastic genomic compartments is an efficient way to regulate the expression of sets of genes scattered throughout the genome that are involved in similar biological processes (Soyer *et al.*, 2015a).

## Conclusion

To the best of our knowledge, previous comparative analyses of histone modifications have been performed in model organisms such as mouse (Mikkelsen *et al.*, 2010) but none considered multiple histone modifications (typical for both heterochromatin and euchromatin) in closely-related phytopathogenic fungi. Nothing is known about how differences in the location of these modifications influence pathogenesis. Comparative genomics has allowed analyses regarding the evolution of genes (notably proteinaceous effectors), the location of effector genes in genomes, or the diversity of effector repertoires in relation to host specialisation or fungal lifestyles. The role of the epigenome is increasingly recognized in plant-pathogenic fungi as an important regulator of genome structure (e.g. Basenko *et al*., 2015; Möller *et al*., 2019; Möller *et al*., 2020) and the expression of genes encoding effectors (e.g. Chujo and Scott, 2014; Soyer *et al*., 2014; Soyer *et al*., 2019). The next step is to exploit comparative epigenomics to better understand the role of the epigenomic landscape in adaptation to environmental changes, modulation of interactions with the holobiont, host adaptation and specialization.

## Author contributions

Conceived, initiated and coordinated the project: JLS, TR, EHS, IF. Designed the experiments: JLS, EHS, IF. Performed the experiments: JLS. Analysis of the data: JLS, CC, EJG, NL. Interpretation of the data: JLS, TR, EHS, IF. Writing of the draft: JLS. Editing of the draft: JLS, TR, EHS, IF.

## Acknowledgments

Authors wish to thank all members of the “Effectors and Pathogenesis of *L. maculans*” and “Environmental Genomics” groups; M. Freitag and J. Grandaubert for fruitful discussions; R. O’Connell for the English proofreading of this paper. J. L. Soyer was funded by a “Contrat Jeune Scientifique” grant from INRA. The “Effectors and Pathogenesis of *L. maculans*” group benefits from the support of Saclay Plant Sciences-SPS (ANR-17-EUR-0007). Research on fungal histone modifications in the group of E.H. Stukenbrock is funded by the Max Planck Society, Germany.

## Supplementary figures legends

**Table S1. Statistics of ChIP-seq and RNA-seq datasets and alignments.** Table shows the number of reads used for the alignment, the number of reads mapping only at one location, the number of reads aligned more than once and the unmapped reads against the genome of *L. maculans* ‘brassicae’ (Lmb; Dutreux *et al*., 2018) or *L. maculans* ‘lepidii’ (Lml; Grandaubert *et al*., 2014).

**Table S2. Correlation analysis between the different ChIP experiments generated with antibodies targeting histone modifications H3K4me2, H3K9me3 and H3K27me3 in *L. maculans* ‘brassicae’, during *in vitro* growth.** Correlation analyses were performed to analyse location of the significantly enriched domains, identified using RSEG (Song and Smith, 2011), from the three different biological replicates generated and analyzed separately. A Kendall’s *𝒯* correlation test was performed using R (Kendall, 1938).

**Table S3. Correlation analysis between the different ChIP experiments generated with antibodies targeting histone modifications H3K4me2, H3K9me3 and H3K27me3 in *L. maculans* ‘lepidii’, during *in vitro* growth.**

Correlation analyses were performed to analyse location of the significantly enriched domains, identified using RSEG (Song and Smith, 2011), from the three different biological replicates generated and analyzed separately. A Kendall’s *𝒯* correlation test was performed using R (Kendall, 1938).

**Table S4. Coverage of histone modifications H3K4me2, H3K9me3 and H3K27me3 in the genome of *Leptosphaeria maculans* ‘brassicae’.**

^a^Genome as published in Dutreux *et al*. (2018)

^b^Location of H3K4me2, H3K9me3 and H3K27me3 was determined through ChIP-seq analysis, *in vitro*, and regions significantly enriched for any of the modifications was identified using RSEG (Song and Smith, 2011).

**Table S5. Coverage of histone modifications H3K4me2, H3K9me3 and H3K27me3 in the genome of *L. maculans* ‘lepidii’.**

^a^Genome as published in Grandaubert *et al*. (2014)

^b^Location of H3K4me2, H3K9me3 and H3K27me3 was determined through ChIP-seq analysis, in vitro, and regions significantly enriched for any of the modifications was identified using RSEG (Song and Smith, 2011).

**Table S6. Correlation analysis between the coverage of transposable elements (TE) and coding sequences (CDS), H3K4me2-, H3K9me3- and H3K27me3-domains in the *L. maculans* ‘brassicae’ genome.** The coverage of transposable elements (TE), coding sequences (CDS) and histone modifications (H3K9me3, H3K27me3, H3K4me2) along the genome of *L. maculans* ‘brassicae’ was analysed in 1000 bp sliding windows. A Kendall’s *𝒯* correlation analysis was done using R.

**Table S7. Correlation analysis between the coverage of transposable elements (TE) and coding sequences (CDS), H3K4me2-, H3K9me3- and H3K27me3-domains in the *L. maculans* ‘lepidii’ genome.** The coverage of transposable elements (TE), coding sequences (CDS) and histone modifications (H3K9me3, H3K27me3, H3K4me2) along the genome of *L. maculans* ‘lepidii’ was analysed in 1000 bp sliding windows. A Kendall’s *𝒯* correlation analysis was done using R.

**Table S8. GO categories enriched in genes associated with H3K4me2 during axenic culture of *L. maculans* ‘brassicae’.** GO annotation of the Lmb genes were retrieved from Dutreux *et al*. (2018). Analysis of GO enrichment among the genes associated with H3K4me2 during axenic culture of Lmb was performed using Cytoscape (Shannon *et al*., 2003).

**Table S9. GO categories enriched in genes associated with H3K4me2 during axenic culture of *L. maculans* ‘lepidii’.** GO annotation of the Lml genes were retrieved from Grandaubert *et al*. (2014). Analysis of GO enrichment among the genes associated with H3K4me2 during axenic culture of Lml was performed using Cytoscape (Shannon *et al*., 2003).

**Table S10. GO categories enriched in genes associated with H3K27me3 during axenic culture of *L. maculans* ‘brassicae’.** GO annotation of the Lmb genes were retrieved from Dutreux *et al*. (2018). Analysis of GO enrichment among the genes associated with H3K27me3 during axenic culture of Lmb was performed using Cytoscape (Shannon *et al*., 2003).

**Table S11. GO categories enriched in genes associated with H3K27me3 during axenic culture of *L. maculans* ‘lepidii’.** GO annotation of the Lml genes were retrieved from Grandaubert *et al*. (2014). Analysis of GO enrichment among the genes associated with H3K27me3 during axenic culture of Lml was performed using Cytoscape (Shannon *et al*., 2003).

**Table S12. Number of (putative) metabolic effector-encoding genes located in different genomic compartments in *L. maculans* ‘brassicae’ and *L. maculans* ‘lepidii’**

^a^Genes located up to 2 kb upstream or downstream of a transposable element sequence;

^b^Genes with RPKM≥ 2;

^c^Genes located in a H3K4me2-, H3K9me3-, H3K27me3- or H3K4me2/H3K27me3-domain *in vitro*; K4: H3K4me2; K9: H3K9me3; K27:H3K27me3.

